# Microtubules coordinate mitochondria transport with myofibril morphogenesis during muscle development

**DOI:** 10.1101/2024.09.16.613277

**Authors:** Jerome Avellaneda, Duarte Candeias, Ana da Rosa Soares, Edgar R. Gomes, Nuno Miguel Luis, Frank Schnorrer

## Abstract

Muscle morphogenesis creates highly specialised muscle cells containing contractile myofibrils and energy producing mitochondria. Myofibrils are chains of sarcomeres, whose myosin motors slide over actin filaments at the expense of ATP. Thus, myofibrils and mitochondria are in intimate contact in mature muscles. However, how mitochondria morphogenesis is coordinated with myofibrillogenesis during development remains largely unknown. Here, we used *in vivo* imaging to investigate myofibril and mitochondria network dynamics in developing *Drosophila* flight muscles. We found that mitochondria rapidly intercalate from the surface of actin bundles to their interior; concomitantly, actin filaments condense to individual myofibrils. This ensures that mitochondria are in intimate proximity to each myofibril. Interestingly, antiparallel microtubules bundle in concert with the assembling myofibrils, suggesting a key role in myofibril orientation. Indeed, light-induced microtubule severing directly affects myofibril orientation, whereas knock-down of kinesin heavy chain specifically blocks mitochondria intercalation and long-range transport. Importantly, mitochondria-myofibril intercalation and microtubule-based transport of mitochondria is conserved in developing mammalian muscle. Together, these data identify a key role for microtubules in coordinating mitochondria and myofibril morphogenesis to build functional muscles.

## Introduction

Complex animals possess several muscle fibre types to support different contraction patterns mediating locomotion and pumping of body fluids. The different fibre types contain particular architectures of their contractile myofibrils and of their energy-producing mitochondria. The most notable difference in mammals is the extremely dense mitochondria network in cardiac muscle, compared to fewer mitochondria in fast glycolytic skeletal muscles, with oxidative skeletal muscles showing an intermediate phenotype^1,2^. This is accompanied by striking biomechanical differences of the sarcomeres, with the rigid cardiac sarcomeres oscillating continuously throughout life, while the more elastic skeletal muscles undergo deliberate boosts of activity that are particularly intense in fast glycolytic fibres and more moderate but long lasting in slow oxidative fibres^3^. These biomechanical differences are largely regulated by differences in the sarcomere protein isoforms^4^, while the metabolism is controlled by their mitochondria content^2^. Thus, muscle physiology depends on the proper coordination of mitochondria morphogenesis with myofibril morphogenesis.

The variability of mitochondria morphologies and their impact on the physiology of different cell types in a complex organism is still underappreciated. Not only the mitochondria density varies in the different muscle types, but also the position and relative orientation of mitochondria with respect to the contractile myofibrils. In oxidative skeletal muscles and particularly in the heart, the mitochondria are located in close proximity to myofibrils^1,5,6^. Hence, mitochondria are maximally close to the ATP-consuming myosin motors.

A similar phenomenon is observed in *Drosophila* adult muscle types. The high energy consuming, oscillating flight muscles contain a particularly high mitochondria density, with mitochondria surrounding each of the individual fibrillar myofibrils, thus supporting them maximally with ATP^7,8^. In contrast, the tubular leg muscles concentrate most mitochondria around their centrally localised nuclei, away from the cross-striated, aligned myofibrils^2,7^. These different architectures are regulated by the master regulator of flight muscle development, the transcriptional regulator Spalt (Salm)^7,9,10^. However, to date we do not understand how the intimate, fibre-type specific interactions between myofibrils and mitochondria are achieved during development.

*Drosophila* flight muscles are an excellent model to study muscle development^11^. Each large flight muscle fibre assembles all of its 2000 myofibrils simultaneously at 30 h after puparium formation (APF), from less well organised actomyosin bundles^12,13^. The assembled myofibrils segregate into large bundles^14^, with mitochondria having intercalated between each of the myofibrils at 32 h APF^7^. Subsequently, myofibrils mature and grow in diameter, while also the neighbouring mitochondria expand in size to fully surround and mechanically insulate the myofibrils at the adult stage^7,15^. How mitochondria-myofibril intercalation and their homogenous distribution along all myofibrils is regulated is not understood.

Skeletal muscle fibres are large syncytial cells that have lost the classical centriole-based microtubule organising centre early during development and instead organise their microtubules around their perinuclear space^16–18^. Microtubules are known to be required for nuclear transport in developing muscles in flies and mammals^19^. They might directly or indirectly also be needed for myofibrillogenesis^20^, though this hypothesis has been challenging to test *in vivo* due to the difficulty of acute microtubule manipulations in such settings. It is well established that mitochondria require microtubules for long-range transport along the axon and dendrites of neurons^21^. However, how mitochondria travel in developing muscles has been little investigated.

Here we found an essential role for microtubules in transporting mitochondria at long-range during the key developmental stage of myofibril assembly in *Drosophila* flight muscles. This microtubule-based transport mechanism is also conserved during mammalian muscle differentiation, ensuring that mitochondria are positioned optimally to fuel the high energy requirements of the myosin motors in mature muscles.

## Results

### Mitochondria intercalate rapidly into actin filament bundles

To study the coordination of myofibril with mitochondria morphogenesis, we used the indirect flight muscles of *Drosophila*. We recently found that mitochondria intercalate between the myofibril bundles between 24 h and 32 h APF^7^. To characterise the dynamics of the intercalation process in detail, we acquired two-colour *in vivo* time-lapse movies, labelling actin and mitochondria, to follow flight muscle morphogenesis in fully intact pupae from 24 h to 30 h APF (Figure 1A, Video S1). These movies revealed interesting dynamics: at 24 h APF, actin filaments are largely located at the periphery of the myotube, while mitochondria are present at distinct central regions (in proximity to the nuclei^22^). At 26 h, actin filaments consolidate into large, distinct bundles throughout the muscle fibre, while mitochondria are located at their surface. Surprisingly, this changes dramatically after 26 h, when most mitochondria intercalate into the actin filament bundles until 28 h APF, while the actin bundles consolidate and form distinct myofibrils at 30 h APF (Figure 1A, Video S1). Thus, mitochondrial intercalation is a rapid process, which appears to be coordinated with myofibril assembly.

**Figure 1:**
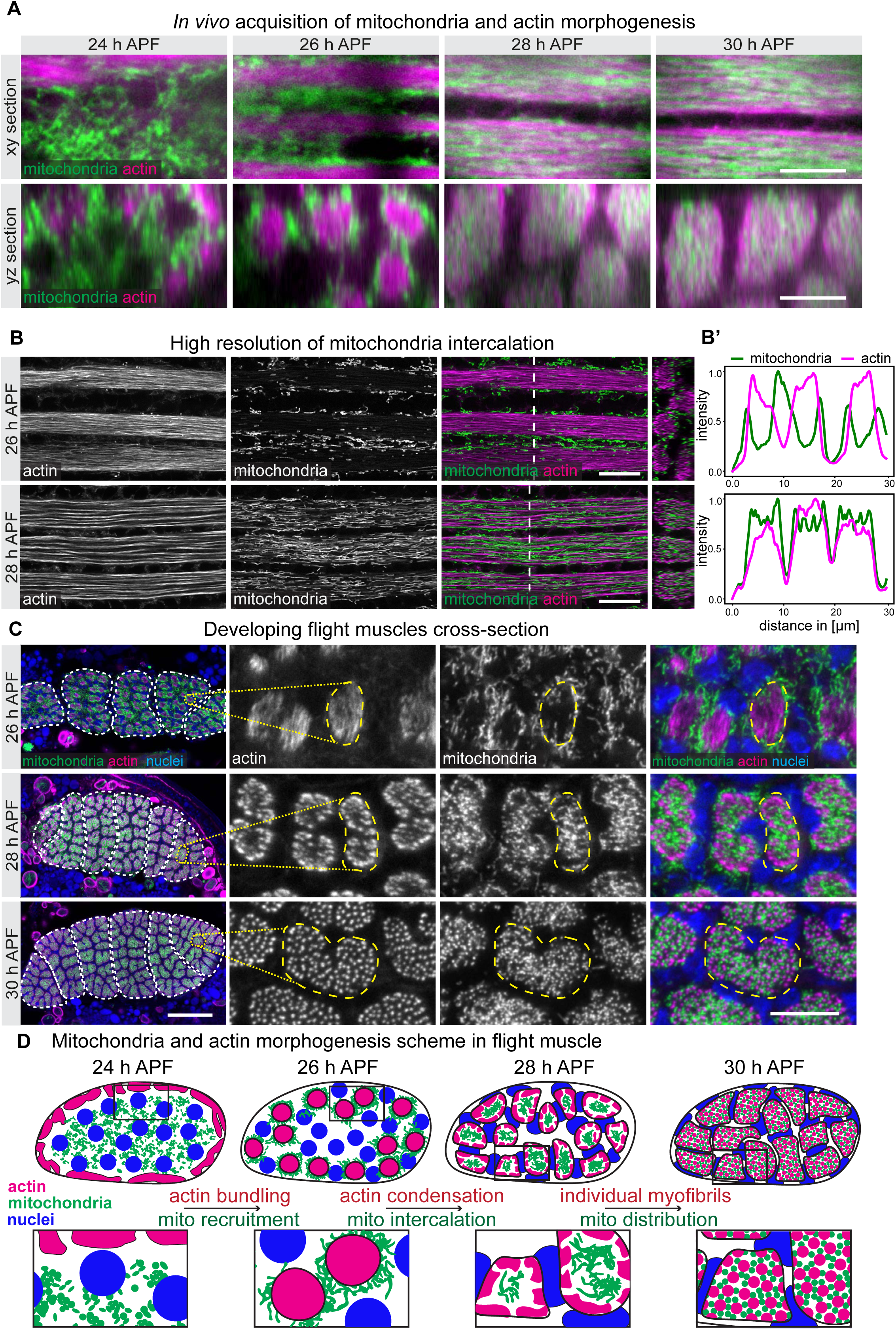
Mitochondria intercalation into actin bundles. **(A)** Stills from a live acquisition of developing flight muscle, labelled for actin with LAD9-GFP and mitochondria with mit-Cherry (expressed with *Mef2*-GAL4) before (24 h, 26 h APF) and after mitochondria intercalation (28 h, 30 h APF); yz cross-sections are shown below. See also Video S1. **(B)** Super-resolution Airyscan images of dissected flight muscles, co-labelled for actin with phalloidin-rhodamine and for mitochondria with mit-GFP enhanced by a GFP nanobody. **(B’)** Intensity profiles of actin (magenta) and mitochondria (green) are plotted along the white dotted line in B; note the overlap of mitochondria and actin at 28 h but not at 26 h APF. **(C)** Cryo-cross-sections of developing flight muscles expressing *Mef2*-GAL4, UAS-mit-GFP at the indicated time points (26 h, 28 h and 30 h APF) were stained with phalloidin-rhodamine (red), GFP nanobody (green) and DAPI (blue) to visualise actin, mitochondria and nuclei, respectively. Left column shows overview cross-sections of 5 - 6 flight muscle cells (white dashed lines); the 3 right-most panels are Airyscan acquisitions of myofibril bundles at high resolution (one bundle is highlighted by the yellow dashed line). Note that mitochondria have intercalated into the bundles at 28 h APF and completely ensheath individual myofibrils at 30 h APF. **(D)** Schemes of mitochondria and actin morphogenesis in flight muscles representing one fiber (top) and a zoom into two actin bundles/myofibril bundles (bottom). The major morphogenesis steps are indicated. Scale bars represent 10 µm in all panels, except the left column C, which represents 50 µm.

To characterise the superposition of mitochondria with actin bundles at high resolution, we performed super-resolution Airyscan confocal imaging of flight muscles fixed at 26 h and 28 h APF. As seen in the live movies, at 26 h APF, distinct actin filament bundles have formed in each muscle fiber. They contain dense actin filaments, which are not yet assembled into individual myofibrils. The mitochondria are excluded from these bundles, while being in close contact at their surface (Figure 1B-B’). At 28 h APF, the actin filaments in each bundle have condensed further generating space in between them, which is now occupied by the intercalated mitochondria (Figure 1B-B’). This demonstrates that mitochondria intercalate into actin bundles concomitantly with the start of myofibrillogenesis.

To observe the relative arrangement of the mitochondria with the assembling myofibrils at even higher resolution, we generated cryo-cross-sections of developing flight muscles. These cross-sections confirmed that mitochondria are in contact with, but largely excluded from, the actin filament bundles at 26 h APF (Figure 1C). Whereas at 28 h APF, actin filaments have condensed at the periphery of each bundle and mitochondria have intercalated into the centre of each bundle. This arrangement becomes more refined at 30 h APF, when all distinct myofibrils have assembled and the mitochondria have redistributed in between them, insulating each myofibril from its neighbours (Figure 1C). Taken together, our results show that mitochondrial reorganisation coincides with myofibrillogenesis. From these observations, we define three main steps: first, dense actin bundles exclude mitochondria at their surface; second, the condensation of actin filaments to myofibrils coincides with mitochondrial intercalation into the bundles; third, when myofibrils have fully assembled, the mitochondria have distributed homogenously between them (Figure 1D).

### Mitochondria dynamics changes upon intercalation

Having shown that mitochondria intercalation is rapid, we aimed to determine the dynamics of mitochondria transport. Expression of GFP fused to a mitochondrial matrix targeting signal (mit-GFP) together with fast frame acquisition (sub-second) determined the individual mitochondria transport speed to be 0,7 µm/s at the pre-intercalation stage at 25 h APF (Figure S1, Video S2). Thus, mitochondria move in the muscle fibres at high speed.

In order to quantify the net translocation of mitochondria throughout the myofiber, we employed the photoconvertible marker Dendra2 localised to the mitochondrial matrix (mit-Dendra2), which allowed us to convert mitochondria from green to red fluorescence in a defined region of the muscle fibre and monitor their transport dynamics (Figure 2A). Surprisingly, by quantifying the speed of translocation along the long axis of the muscle at 25 h APF, we found that mitochondria move in bulk only over relatively short distances within 30 minutes (Figure 2B, F, Video S3). Conversely, after mitochondrial intercalation at 32 h APF, mitochondria translocate rapidly across large distances along the long axis of the flight muscle fibre (Figure 1C, F, Video S3). This demonstrates that mitochondria transport dynamics changes dramatically in the muscle cells, promoting mitochondria long-range transport after intercalation occurred.

**Figure 2:**
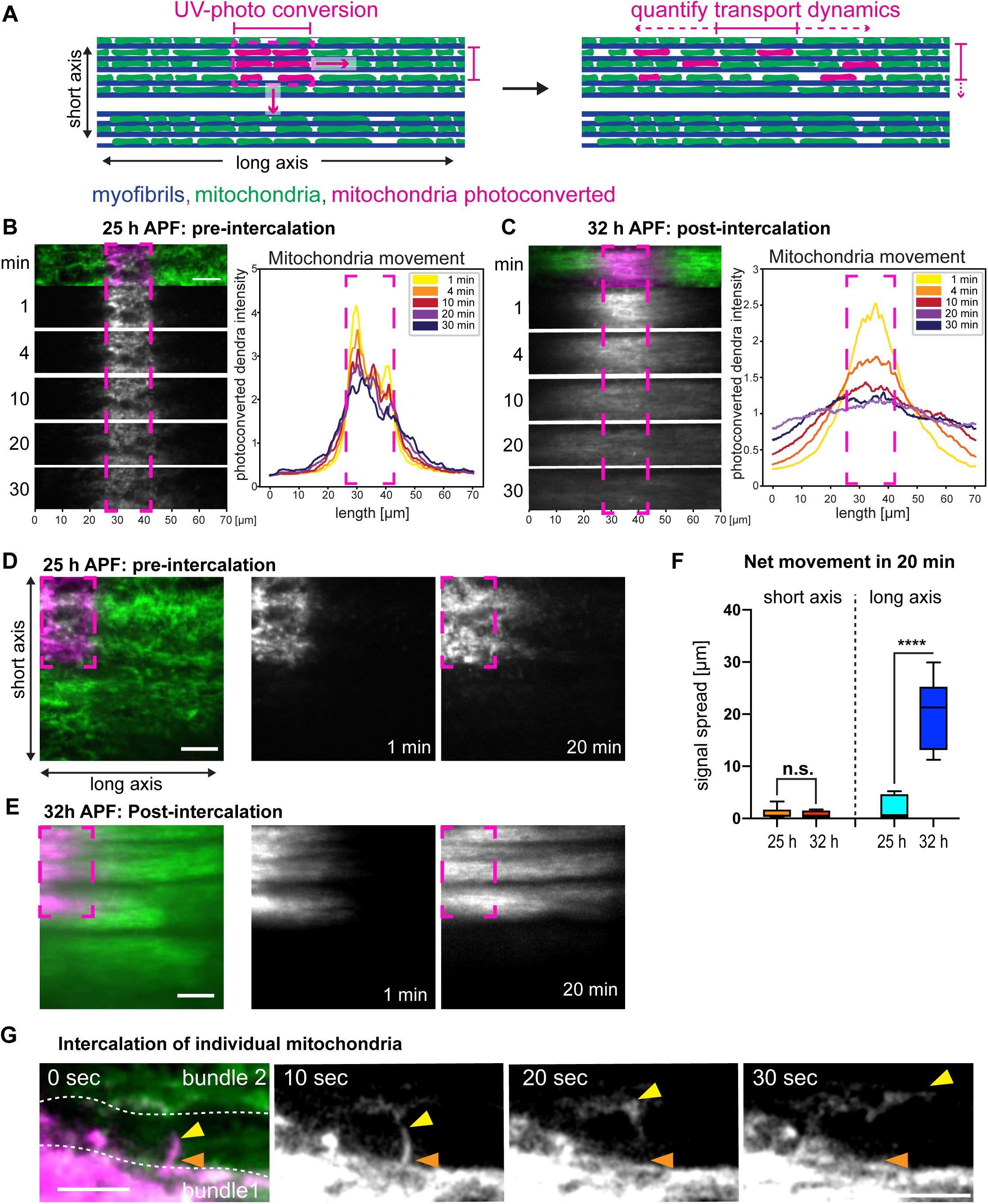
Mitochondria transport dynamics switches after intercalation. **(A)** Schematic overview of a photo-conversion experiment using mit-Dendra2 and *Mef2*-GAL4 to label mitochondria in flight muscles. (**B, C)** Time lapse live imaging of developing flight muscles expressing mit-Dendra2 (driven by *Mef2-*GAL4) at 25 h APF (B) and 32 h APF (C), after photo-conversion. The magenta dashed rectangles label the photo-converted area. Top images show a composite of non-converted mit-Dendra2 in green and photo-converted mit-Dendra2 in magenta. The photo-converted mit-Dendra2 is shown in grey below at the indicated time after conversion. Plot profiles quantify the photo-converted mit-Dendra2 intensities over time. Note the slow spread at 25 h APF compared to the fast spread at 32 h APF. See also Video S3. **(D, E)** Photo-conversion of mitochondria at only a thin rectangular area of a muscle fiber allows to compare the mit-Dendra2 spread along the short- and long-axis of the cell before intercalation at 25 h APF (D) or after intercalation at 32 h APF (E). Images were acquired immediately after (1 min) or 20 minutes after photoconversion. See also Video S4. (**F**) Quantification of the fluorescence signal spread after 20 min along the short or long axis at 25 h and 32 h APF from the acquisitions shown in D and E. Note the virtually absent spread along the short axis. **(G)** Single frames from a live acquisition (see Video S5) in which a single photo-converted mitochondrion intercalates into an actin bundle within one minute (orange arrowheads mark the start at 0 sec, the yellow arrowheads the tip of the mitochondrion during the movie). Significance from two-tailed unpaired *t*-tests \*\*\*\**p* ≤ 0.0001, (n.s.) non-significant. All scale bars represent 10 µm.

Having identified this fast transport dynamics, we wanted to clarify how directed the movement is. To this end, we photo-converted a smaller region per muscle fibre and followed the translocation of mitochondria along the long and the short axis of the muscle. At 25 h APF, we found almost no translocation along either axis (Figure 2D, F, Video S4). At 32 h APF, we confirmed the above identified fast translocation of the mitochondria along the long axis of the muscle. However, along the short axis, we found no translocation (Figure 2E, F, Video S4). These results show that the long-range mitochondria translocation is specifically oriented along the long axis, strongly suggesting an active transport mechanism.

### During intercalation mitochondria move along the short axis into actin bundles

Having shown that mitochondria gain long-range movement after intercalation, we wanted to follow them directly during the intercalation process. We selected the 27 h APF time point, while mitochondria intercalation is ongoing, photo-converted only a few mitochondria and followed them at high frame rate. Using this setup, we found that mitochondria can move also along the short axis into the actin filament bundles in less than 1 minute (Figure 2G, Video S5). This suggests that also the short axis movement of mitochondria during mitochondria-myofibril intercalation is likely using an active transport mechanism.

### Actin and microtubules bundles are closely associated

How are mitochondria transported during flight muscle development? The microtubule cytoskeleton is the obvious candidate, as it is the main transport route for mitochondria in various cell types^23^; however, the function of microtubules is not well explored during flight muscle development^20^. We first analysed the relation of microtubules to the actin filaments during flight muscle development. By imaging flight muscles expressing Lifeact-Ruby and Tau-GFP to mark actin and microtubules, respectively, we found that both co-localise extensively (Figure S2A, Video S6). At 24 h APF, actin filaments and microtubules largely accumulate at the periphery of the myotube, while a microtubule mesh is visible in the centre, around the centrally located nuclei (dark circles^22^). At 26 h APF, actin filaments and microtubules start to bundle together simultaneously and as myofibrils assemble at 29 h APF, microtubules and actin remain closely associated (Figure S2A, Video S6).

To increase the resolution, we generated longitudinal optical sections and physical cross-sections of flight muscles expressing β-Tub60D-GFP to label the microtubules that were co-stained with phalloidin between 26 h and 30 h APF and imaged with Airyscan super-resolution microscopy (Figure 3A, B). This confirmed the high concentration of the microtubules in the assembled actin filament bundles at 26 h APF, with some microtubules still localised between actin bundles. This is consistent with mitochondria being transported along microtubules into the actin filament bundles after 26 h APF. This inter-bundle microtubule mesh is still observed at 28 h APF, while it disappears at 30 h APF (Figure 3A, B). At 30 h APF, we found the microtubules organised around each individual myofibril (Figure 3B, B’), which is consistent with electron-microscopy data of the same stage^24,25^. The space between the microtubules is probably occupied by the intercalated mitochondria at 30 h APF (Figure 3B, B’). Microtubules remain around the myofibrils until 48 h, while they disappear when myofibrils have grown in diameter at 72 h or 90 h APF (Figure S2B). Taken together, our results show a close association between the actin and the microtubule cytoskeleton before and during myofibrillogenesis, which supports a possible role of microtubules during mitochondria intercalation.

**Figure 3:**
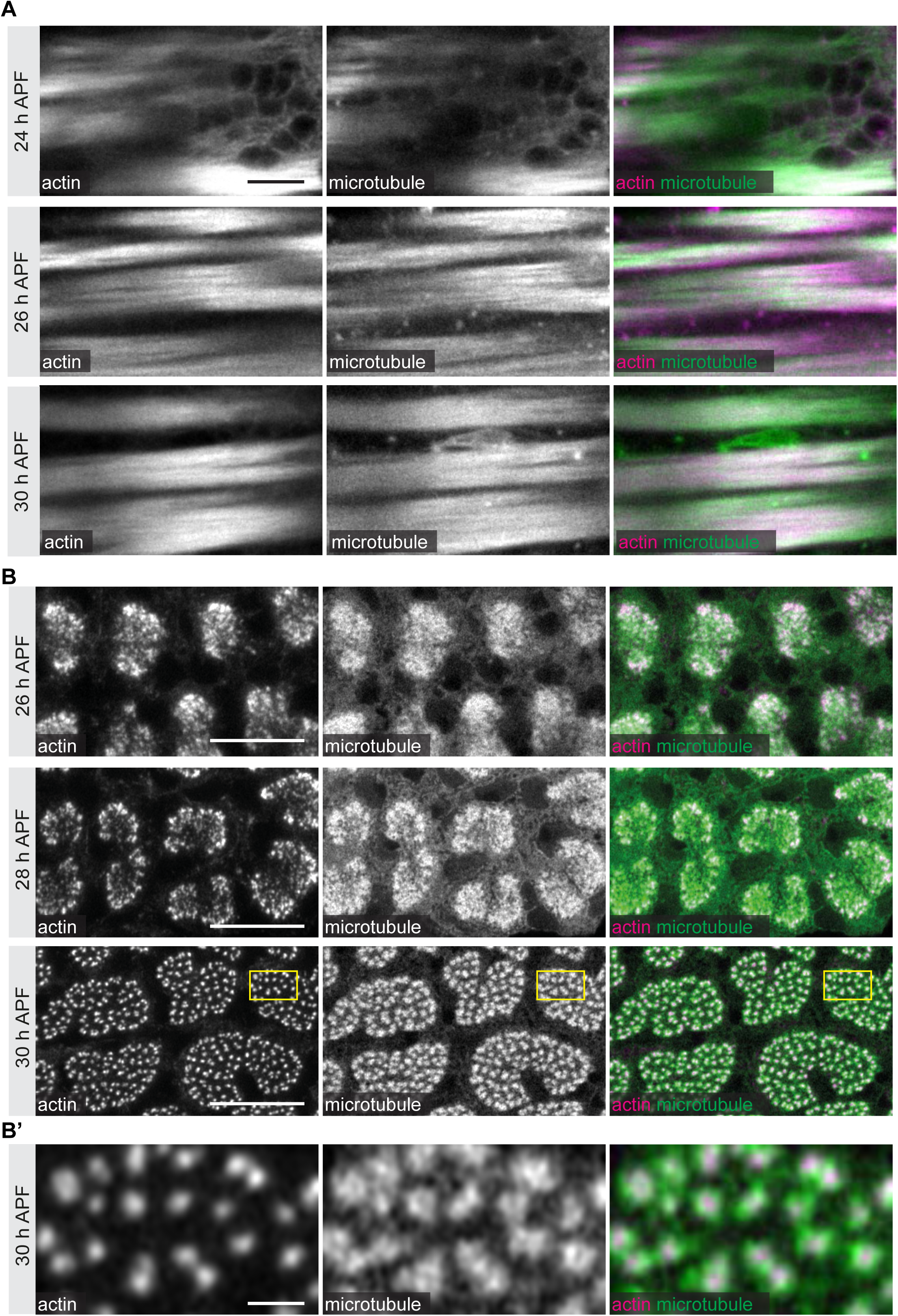
Myofibrils form in close association with microtubules. **(A)** Longitudinal confocal images of developing flight muscle at the indicated time points labelled for actin (Lifeact-Ruby, driven by *Mef2-*GAL4) and for microtubules (*Mhc*-Tau-GFP). Note how actin and microtubules overlap at this resolution. See also Video S6. **(B)** Airyscan high-resolution acquisitions of flight muscle cross-sections at indicated time points (26 h, 28 h and 30 h APF) labelled for actin (phalloidin) and microtubules (β-Tub60D-GFP). **(B’)** Zoom-in of the 30 h APF stage at the indicated yellow box in B. Note that microtubules largely surround each assembled myofibril at this stage. Scale bars represent 10 µm in A and B, and 1 µm in B’.

### Mitochondria follow microtubules into the actin bundles

To visualise microtubules and mitochondria simultaneously, we co-labelled them in developing flight muscles by expressing β-Tub60D-GFP and mit-Kate2. We found that mitochondria are excluded from the microtubule bundles at 26 h APF, whereas they have intercalated inside the bundles at 28 h APF (Figure 4A). This is consistent with our above findings that actin filament bundles intimately associate with microtubules (Figure 3A, B) and that mitochondria intercalate into the actin filament bundles (Figure 1B). Importantly, the few non-intercalated mitochondria, that are still present outside the microtubule bundles, are often associated along transversely oriented microtubules that can mediate transport along the short muscle axis (Figure 4B). This microtubule population probably corresponds to the mesh of microtubules best observed in cross-sections (Figure 3B). Overall, these results support a mechanism by which mitochondria intercalate into the actin filament bundles through microtubule-dependent transport.

**Figure 4:**
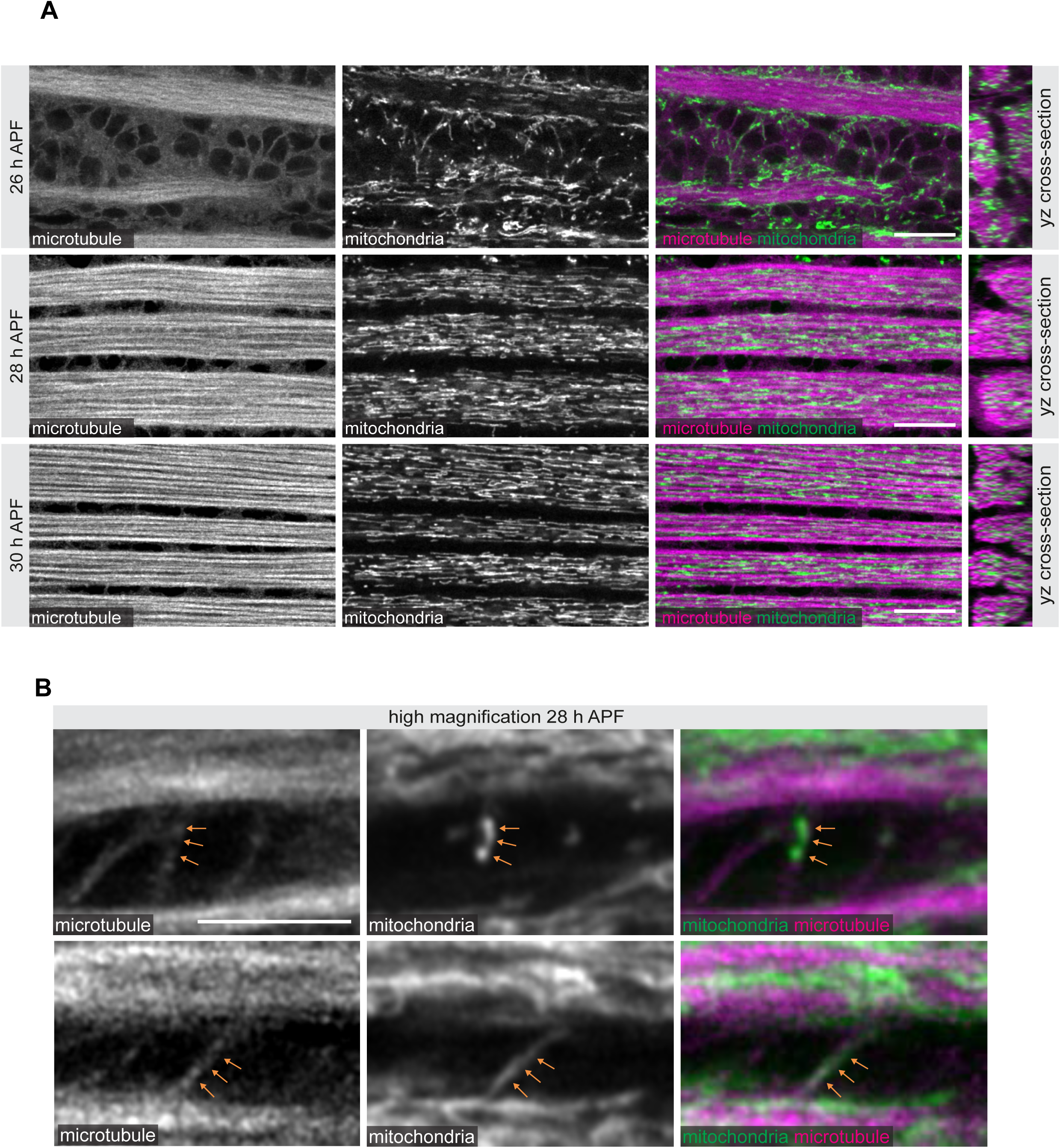
Mitochondria are associated with microtubules. **(A)** Flight muscles labelled for mitochondria (mit-Kate2, expressed via *Mef2-*GAL4) and microtubules (*β-Tub60D*-GFP) at the indicated developmental time points (26 h, 28 h and 30 h APF) and imaged with Airyscan super-resolution microscopy. Note that mitochondria are located on the surface of microtubule bundles at 26 h APF, while they have intercalated into these bundles at 28 h APF. yz optical cross-sections are shown on the right. **(B)** Close-ups of similar images shown in A focussing on two different single mitochondria that are not yet intercalated (labelled by orange arrows; top and bottom). Note their close association with microtubule filaments at 28 h APF. Scale bars represent 10 µm (A) and 5 µm (B).

### Microtubules are required for actin bundling and mitochondria positioning

To functionally test the microtubule-dependent mitochondrial intercalation hypothesis, we aimed to deplete microtubules by expression of the microtubule severing protein Spastin^26–28^. However, since microtubules are important for many cellular functions, the expression of Spastin must be precisely controlled in time. To do so, we employed the recently developed photoactivatable *shine-* GAL4, which enables light-controlled expression within 1-2 hours^29^. *shine-*GAL4 UAS-*spastin* flies raised in the dark are wild type, while exposing them to light starting at 23 h APF is pupal lethal. This demonstrates the absence of leakage of *shine*-GAL4 activity in the dark.

To test the effect of Spastin expression on myofibrillogenesis, we raised *shine*-GAL4 UAS-*spastin* pupae in the dark and exposed them to light at 23 h APF, and then analysed the flight muscles at 32 h APF (Figure 5A, B). We found that these flight muscles show severe myofibril defects: myofibrils do not form distinct bundles and run in variable directions in the same muscle fibre, compared to their regular arrangement in bundles in control pupae raised in constant darkness (Figure 5B). Furthermore, the nuclei are not restricted to the inter-bundle space as they are in wild-type fibres but distributed more homogenously (Figure 5B). This demonstrates that microtubules are required after 23 h APF for myofibrillogenesis and correct myofibril orientation. To directly investigate a role of microtubules for mitochondria transport and actin filament bundling, we exposed *shine*-GAL4 UAS-*spastin* pupae to light at 23 h APF and analysed their flight muscles only 2 hours later at 25 h APF (Figure A, C). We found that microtubule severing during this brief developmental period is sufficient to completely block actin filament bundle formation. Instead, actin filaments form a homogenous mesh with interspersed nuclei, and consequently the mitochondria are not re-organising into their bundle-associated pattern as they do in controls (Figure 5C). This demonstrates that microtubules are critical for actin filament bundling and mitochondria localisation.

**Figure 5:**
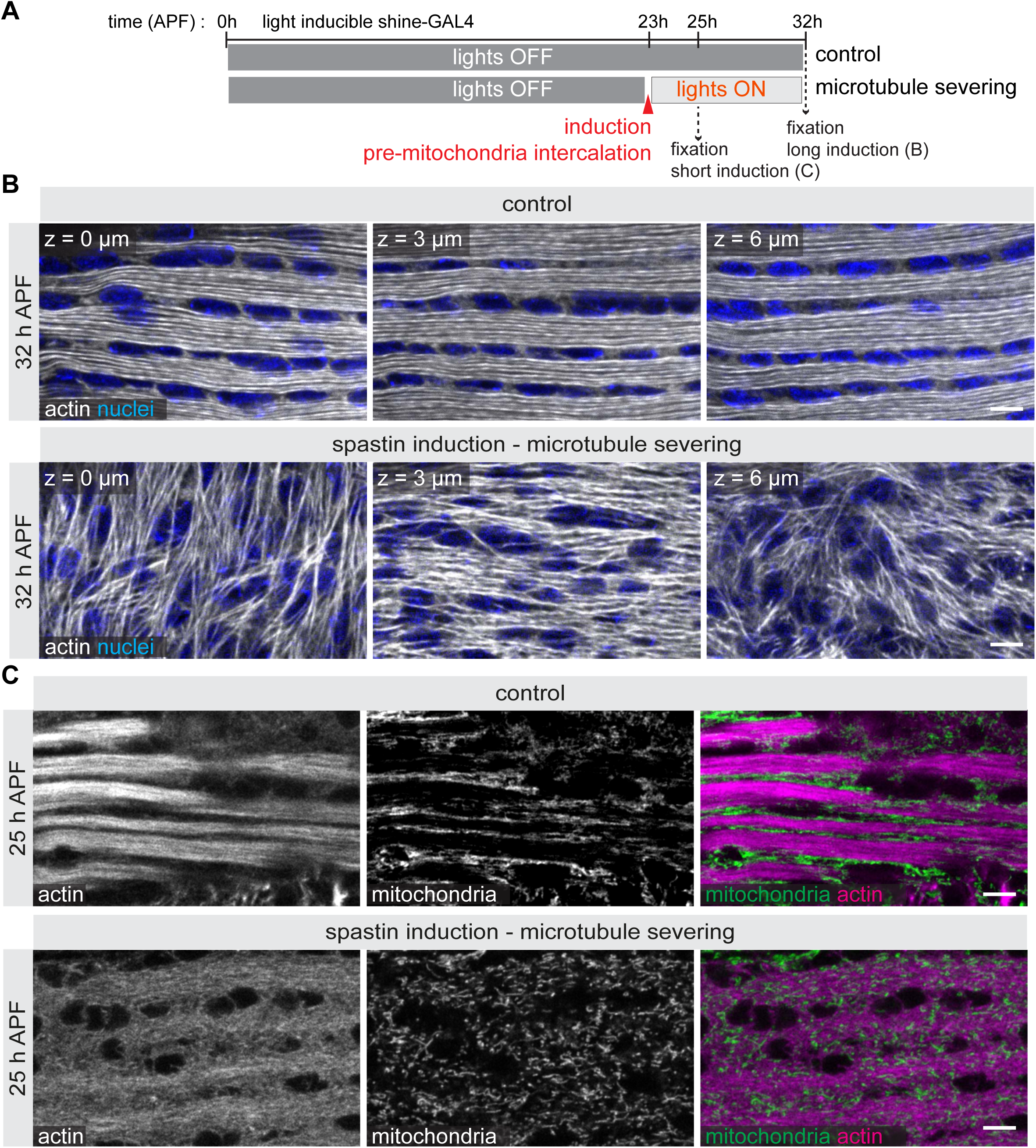
Microtubules are required for myofibrillogenesis. **(A)** Scheme of temporal Spastin expression via light-activation with the shineGAL4 driver and the indicated sample collection time points. **(B, C)** Confocal images of developing flight muscles from shine-GAL4; UAS*-Spastin* pupae raised in the dark (control, top rows in B and C) or exposed to light after 23 h APF to induce Spastin expression (bottom rows in B and C) and labelled for actin (phalloidin) and nuclei (DAPI) at 32 h APF (B) or labelled for mitochondria (ATP5α antibody) and actin (phalloidin) at 25 h APF (C). Note the severe myofibril orientation defects upon Spastin expression in (B) and the defects in actin bundling in (C). Scale bars represent 5 µm.

Inducing the expression of Spastin only after 30 h APF does not strongly perturb mitochondria positioning relative to the myofibrils at 48 h, while mitochondria appear slightly smaller and sarcomeres slightly shorter at 90 h APF (Figure S3). This suggests that microtubules have no major role in myofibril maturation, whereas they are needed during the preceding step of actin bundling, myofibril assembly and mitochondria intercalation.

### Kinesin is necessary for mitochondria intercalation

As the complete disassembly of microtubules affects the bundling of the actin filaments, it is trivial that the mitochondria are also mis-localised. To investigate more specifically the role of microtubule transport in mitochondria intercalation, we have performed a candidate RNAi-screen and identified an important role for the microtubule motor *kinesin heavy chain* (*khc*). Knocking-down *khc* in muscle by RNAi (*khc-IR*) using *Mef2*-GAL4 results in late pupal lethality, demonstrating an important role for *khc* for muscle development or muscle function^30^. Investigating the flight muscle morphology of these late pupae showed a dramatic agglomeration of mitochondria outside of the myofibril bundles in mature muscles at 90 h APF, demonstrating that *khc* is indeed important for normal mitochondrial localisation (Figure S4A, B). As a consequence, the diameter of the *khc-IR* myofibrils is enlarged and more variable compared to wild type (Figure S4C, D). This suggests that kinesin is the motor protein transporting mitochondria during mitochondria intercalation. To directly test this, we analysed *khc-IR* flight muscles at 32 h APF and found that actin filaments bundle normally, with myofibrils assembling. This suggests that the microtubule network reorganises together with the actin filaments and remains intact upon *khc* knock-down. However, most mitochondria remain in the space outside of the myofibril bundles in large aggregates (Figure 6A, B). Thus, kinesin is required for mitochondria intercalation into the myofibril bundles before 30 h APF.

**Figure 6:**
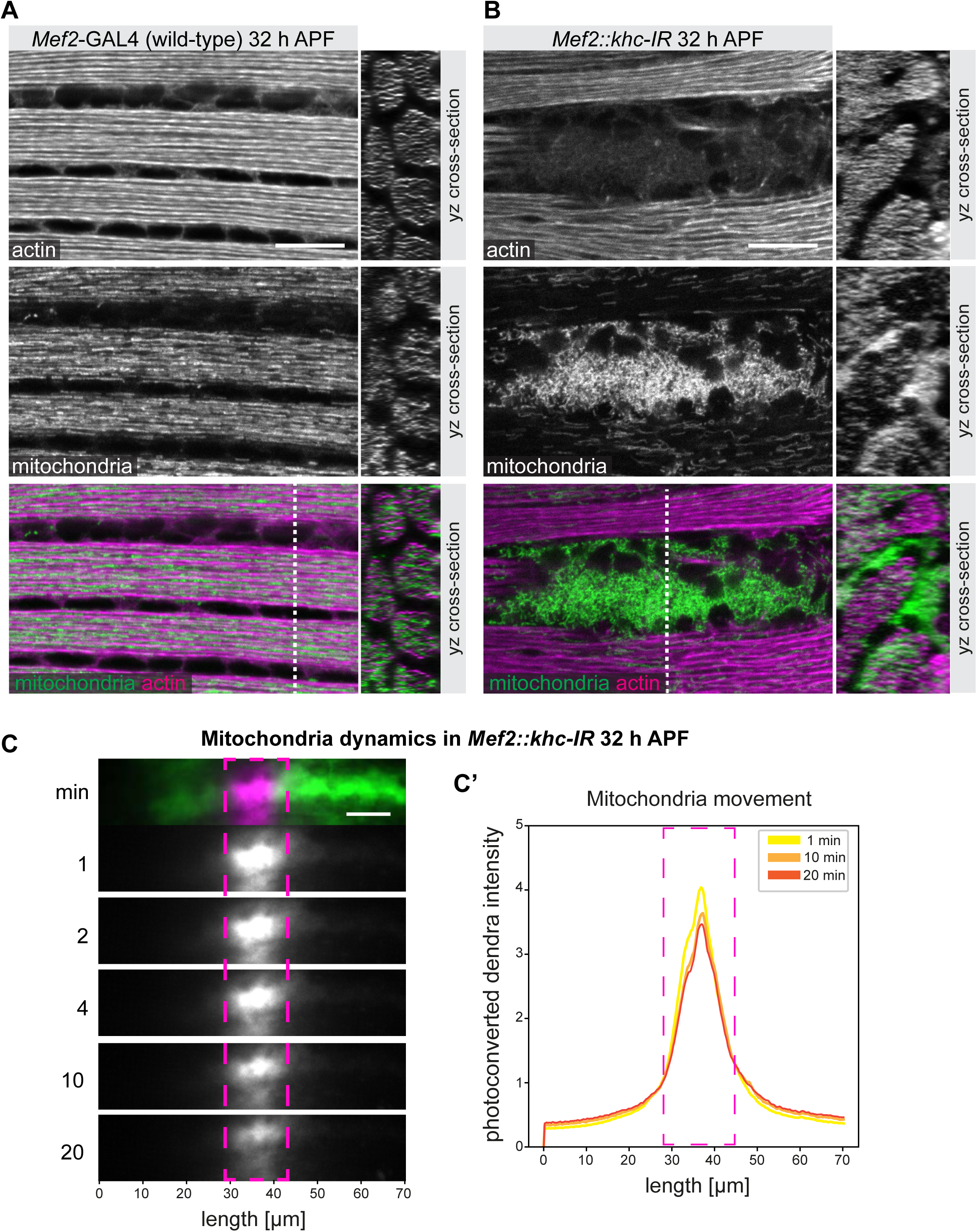
Kinesin dependent mitochondria transport. **(A, B)** Flight muscles from control (*Mef2-*GAL4) (A) or kinesin knockdown (*Mef2::khc-IR*) (B) pupae were labelled at 32 h APF for actin (phalloidin) and mitochondria (mit-GFP). x-y views are shown to the left and y-z sections at the indicated white dotted lines to the right. Note how kinesin knock-down results in large mitochondria clusters and blocks intercalation. **(C, C’)** Time lapse live imaging of mit-Dendra2 in developing flight muscles at 32 h APF after kinesin knockdown (*Mef2::khc-IR*). The magenta dashed rectangle labels the photo-converted area. Top row shows a composite of non-converted mit-Dendra2 in green and photo-converted mit-Dendra2 in magenta. The photo-converted mit-Dendra2 is shown in grey at the indicated time after conversion. Plot profile (C’) quantification of the photo-converted mit-Dendra2 intensities over time. Note the nearly complete absence of spread compared to the fast spread at 32 h APF in control muscle in Figure 2C. All scale bars represent 10 µm.

Finally, we tested the mobility of the mitochondria in *khc-IR* muscles using our established photoconversion assay of labelled mitochondria. We found that in contrast to wild type, mitochondria in *khc-IR* flight muscles remain immobile at 32 h APF (Figure 6C). This demonstrates that kinesin is required to transport mitochondria at long range in developing flight muscles during and after intercalation.

### Mitochondria intercalation and microtubule dependent transport is conserved in mammalian muscles

To test if the microtubule-based mitochondria reorganisation mechanism is conserved in developing mammalian muscles, we investigated mitochondria localisation in mouse primary myofibers differentiated *in vitro*, which recapitulate most stages of early muscle morphogenesis^31^. According to actin filament morphologies and nuclear positioning, we classified developing myofibers into one of four types (Figure 7A, Figure S5): in type 1, mitochondria are largely excluded from a dense actin filament mesh at the cell cortex and rather cluster around the nuclei. In types 2 and 3, nuclei have distributed and actin filaments have oriented along the fibre axis, with the first periodic myofibrils appearing, while mitochondria progressively locate in between the assembling myofibrils. Finally in type 4, the nuclei have moved to the periphery, the sarcomeres have matured into periodic repeats and neighbouring myofibrils have aligned into their typical cross-striated pattern. Interestingly, the mitochondria are located in between bundles of myofibrils and appear elongated along the long axis of the type 4 fibres, while some mitochondria are even present at the periphery of the type 4 fibre (Figure 7A, Figure S5). The transition from type 1 and 2 myotubes to type 3 and 4 myofibers takes about 4 days in culture (Figure 7A, B). This is consistent with previous observations and verified our culture conditions^31^. This shows that within 4 days of culture, mouse mitochondria dynamically change their location, concomitantly with myofibrillogenesis, very reminiscent of the intercalation observed in the developing insect flight muscle.

**Figure 7:**
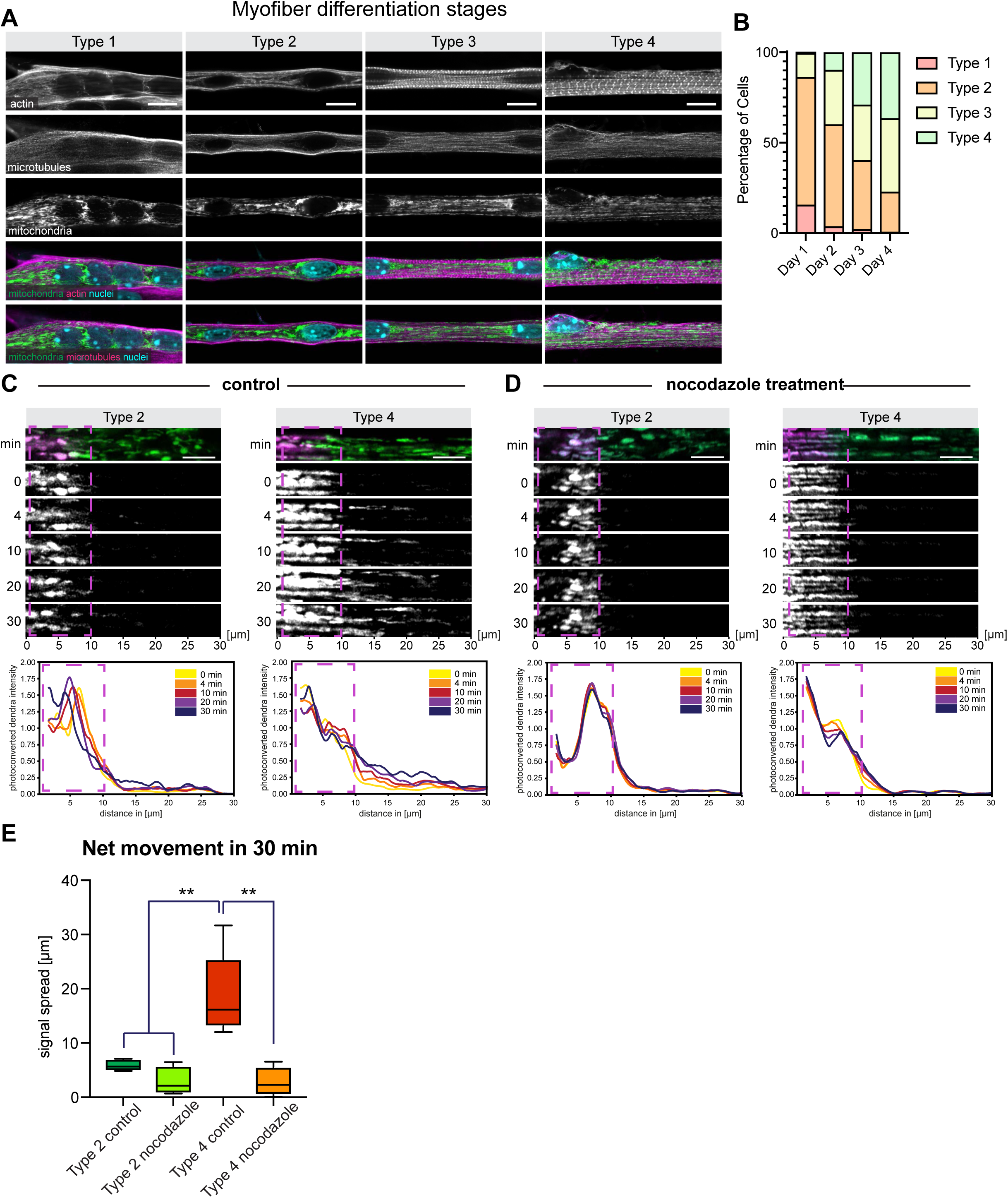
Microtubule-dependent transport of mitochondria is conserved in mammalian muscle. **(A)** *in vitro* differentiation of mouse myoblasts into mature myofibres classified into 4 differentiation types. Cells were transfected with 4xmts-mNeonGreen to label mitochondria, fixed and labelled with phalloidin, anti-tyrosinated α-tubulin (YL1/2), and DAPI, to label actin, microtubules and nuclei, respectively. Criteria for classification are shown in Figure S5. **(B)** Quantification of the myofibre type distribution during 4 days of culture shows a progressive increase in differentiated myofibers (type 3 and 4). **(C, D)** Cultured myofibres were transfected with mit-Dendra2 and mitochondria were photo-converted at the indicated magenta boxes. Type 2 and type 4 fibres were imaged immediately after photoconversion and then every 2 min. Top images show a composite of non-converted mit-Dendra2 in green and photo-converted mit-Dendra2 in magenta. The photo-converted mit-Dendra2 is shown in grey below at the indicated time after conversion. In (D) the cultured myofibers were treated with 10 µg/mL nocodazole, 5 min prior to photo-conversion. Plot profiles below quantify the photo-converted mit-Dendra2 intensities over time. Note the slow spread of type 2 compared to the faster spread of type 4 myofibres, and the absence of spread upon nocodazole treatment. Scale bars are 10 µm in (A) and 5 µm in (C) and (D). **(E)** Quantification of the fluorescence signal spread after 30 min from the acquisitions shown in (C) and (D). Significance from two-tailed unpaired *t*-tests \*\**p* ≤ 0.01, scale bars represent 10 µm in A and 5 µm in B, C.

To test how mitochondria dynamics changes during myofiber morphogenesis, we employed our above-described mitochondria photoconversion experiment by transfecting myotubes with mit-Dendra2. We photoconverted mitochondria in muscle cells of type 2 and type 4 (before and after the mitochondria re-locate, respectively) and followed the spread of the signal to quantify mitochondria long-range movement. Similar to the pre-intercalation stage in flight muscles, we found little mitochondria long-range translocation at stage 2, while at stage 4 mitochondria move over longer distances, similarly to the post-intercalation stage in the flight muscle (Figure 7C).

To directly test for a function of the microtubules to instruct mitochondria dynamics we acutely depolymerised the microtubules with nocodazole and imaged mitochondria after photoconversion, 5 min after treatment with nocodazole. We found mitochondria long-range transport stops almost completely after nocodazole treatment (Figure 7D, E).

Taken together, these data demonstrate that mitochondria are transported along the microtubule cytoskeleton, resulting in a dramatic change of their location, in order to coordinate with the assembling and maturing myofibrils during mouse myofiber differentiation. Hence microtubules coordinate mitochondria transport with myofibril morphogenesis in developing mouse myofibers.

## Discussion

The two key components of muscles are the contractile myofibrils and the energy producing mitochondria. Depending on the biomechanical and physiological properties of the muscle type, the interplay between these components results in fibre-type specific relative arrangements^1,5,12^, which are important for proper muscle function. In the *Drosophila* flight muscle, mitochondria intercalate between the assembled myofibrils during development; this intercalation then even feedbacks on the transcriptional program of the muscle type^7^. Hence, the morphogenesis of mitochondria and myofibrils must be tightly coordinated.

Here we investigated the mechanism how mitochondria intercalate in between the developing myofibrils in flight muscles. We found that this intercalation is a rapid active process: the mitochondria, initially located centrally in the myotube close to the nuclei, intercalate into the actin filament bundles within two hours. Interestingly, this intercalation coincides precisely with the time of myofibril assembly: at 30 h APF, periodic immature sarcomeres are present in all of the 2000 myofibrils that simultaneously assemble in each flight muscle fibre^13,32^. Thus, the intercalation between these assembled myofibrils happens homogenously throughout the cell, resulting in 2000 myofibrils, each of which is surrounded by long tubular mitochondria^24,25^. We revealed a very similar simultaneous change in mitochondria localisation during assembly of the cross-striated myofibrils in developing mouse muscle fibers.

We identified the mechanism how the rapid intercalation of mitochondria is achieved. We discovered an active transport mechanism of the mitochondria into the myofibril bundles by the microtubule cytoskeleton. The evidence for this transport mechanism is fivefold: first, we found that microtubules are very closely associated with the actin filaments or the assembling myofibrils during the stage of mitochondria intercalation. Second, we documented a rapid long-range transport of the mitochondria along the orientation of the microtubule axis, after intercalation, both in flight and mammalian muscles. Third, we showed that severing microtubules blocks actin bundle formation and mitochondria redistribution. Fourth, we identified the kinesin motor protein as essential for mitochondria intercalation into the myofibril bundles. Finally, we showed directly that kinesin is required for long-range mitochondria transport after myofibril assembly. Hence, these observations provide very strong evidence for our hypothesis that the microtubule cytoskeleton directs the mitochondria into the proximity of individual myofibrils both in flight as well as in mammalian muscles. The high longitudinal translocation speed and the anti-parallel microtubule organisation ensures an equal distribution of the mitochondria within each of the myofibril bundles: as mitochondria continuously move back and forth along the myofibril axis of the developing muscles, they stay distributed.

How is the transport initiated and how is kinesin recognising the mitochondria to be transported? The standard kinesin-mitochondria adaptors Milton and Miro, which are required for transport in neurons^33,34^, do not seem to be involved, as knock-down of either Milton or Miro does not show a phenotype in the flight muscle^30^. Which other adaptors could be involved is unclear to date. Further, we do not know why the intercalation is so rapid and precisely staged, and not already occurring before 26 h APF, when a large number of microtubules are already present within the actin filament bundles. A plausible mechanism could be that the early actin bundles are forming a branched mesh with a small pore size, and hence the rather large mitochondria cannot intercalate and remain at the bundle surface. When the actin filaments in each bundle condense to defined myofibrils, enough space between the individual myofibrils is generated, allowing the kinesin motors to drag mitochondria in between the assembling myofibrils. This hypothesis is supported by the observation that induced hyper-fusion resulting in larger mitochondria, blocks the intercalation entirely^7^.

What is the physiological advantage for muscles to apply such an intercalation mechanism? We see at least three advantages. First, it ensures that each myofibril is surrounded by a maximal number of mitochondria. As these will grow in volume when development proceeds, each mature sarcomere is in intimate proximity to the ATP producing source. Second, the rapid transport along the microtubules next to the assembled myofibrils ensures that mitochondria are equally distributed around all of the 2000 myofibrils per flight muscle cell and along the cross-striated myofibrils of mammalian fibres. If mitochondria would stay clustered, only the ones close to the cluster would be supported with ATP. Third and most important, the intercalation allows to limit the diameter growth of the myofibrils by physically blocking the space in between them until a maximum pressure is reached^7,35^. Thus, the sarcomeres can maintain an optimal diameter that is compatible with maximal ATP support by the proximal mitochondria. This is supported by our finding that the myofibril diameter in flight muscles increases and displays a larger variation when mitochondria intercalation was blocked.

The proposed transport mechanism requires that every individual myofibril is surrounded by several microtubules extending along its length. This has been found to be the case at 32 h APF using high-resolution electron microscopy^24,25^. Our super resolution imaging of flight muscle cross-sections and developing mammalian muscles also showed that microtubules progressively surround the nascent myofibrils, as actin filaments condense to highly organised myofibrils. Once mitochondria are inside the bundles, the microtubules provide a highway for the transport and homogenous distribution of mitochondria along a developing and growing muscle.

How do microtubules organise themselves in proximity to the actin filament bundles? At the moment, we can only speculate. It is well established that microtubules are organised around the many nuclei of the muscle fibres^18,36^. However, at the intercalation stage the nuclei are located outside of the myofibril bundles, at the surface of mature mammalian fibers, and in between the actin bundles in flight muscles, thus not where most microtubules are found. Hence, it is more likely that the assembling myofibrils recruit or organise the microtubules and vice versa, as supported by the dramatic actin filament defects found within 2 hours after microtubule severing. One candidate for such linker protein between actin and microtubules is Shortstop (spektraplakin)^37–39^. Microtubules could then act as an orientation cue and help actin polymerisation along the long axis of the muscle as has been suggested^40^. Thus, microtubules first organise the actin cytoskeleton into myofibrils and then ensure that each myofibril will be surrounded by the energy producing mitochondria.

The detailed arrangement will be adjusted to the muscle type. As the mammalian myofibrils are cross-striated, the ensheathing of each myofibril by the mitochondria is less complete compared to the flight muscle. Some mitochondria only send protrusions to the mature mammalian sarcomeres at defined positions of the sarcomere^1^. A similar observation was made in cross-striated *Drosophila* leg muscles, with their mitochondria projecting thin extensions to the sarcomere Z-discs^2,7^. If these particular mitochondria patterns also depend on microtubule transport or anchoring will need further investigations.

## Materials and Methods

### Fly strains and genetics

All crosses were carried out at 27°C under standard conditions^7^. Flight muscle development movies used *Mef2*-GAL4, UAS-mit-Kate2; UAS-LAD9GFP (affimer binding actin, gift of Manos Mavrakis^41^) pupae for double labelling of mitochondria and actin. *In vivo* double labelling of mitochondria and microtubules was performed with *Mef2*-GAL4, UAS-mit-Kate2 ^15^; *Mhc-* TauGFP (BL53739)^42^ pupae. *In vivo* double labelling of actin and microtubules was performed with *Mef2*-GAL4, UAS-LifeAct-Ruby (BL35545)^43^; *Mhc*-TauGFP pupae. Photoconversion experiments used *Mef2*-GAL4, UAS-mit-Dendra2 ^44^ pupae. Spastin induction experiments were performed with shine-GAL4 ^29^; UAS-*spastin-*EGFP ^26^. Experiments using fixed flight muscles were performed with *Mef2*-GAL4; UAS-mit-GFP (BDSC: 25747) for Figure 1 or *βTub60D*-GFP fosmid flies^45^ for Figure 3 or *Mef2*-GAL4, UAS-mit-Kate2; *βTub60D*-GFP flies for Figure 4. Kinesin knockdown experiments were performed with *Mef2*-GAL4 crossed to *w[1118] control* or UAS*-khc-IR* (VDRC: 44337, GD12278)^46^.

### Cell culture of primary myofibers

All procedures using animals were approved by the institutional ethics committee and followed the guidelines of the National Research Council Guide for the care and use of laboratory animals. *In vitro* myofibers were differentiated as previously described^47^ from primary myoblasts isolated from dorsal hind-limb muscles of 5–7-day-old C57BL/6 mice. Briefly, muscles were dissected, minced mechanically with curved scissors and digested with collagenase type V (Sigma-Aldrich #C9263) and dispase II (Gibco #17105-041) in DPBS (Gibco #14040133) for 90 min at 37°C with agitation. The digestion was stopped by mixing IMDM with Glutamax (Gibco #31980022), 10% FBS (Eurobio Scientific #CVFSVF00-01) and 1% Penicillin-Streptomycin (Gibco #15140-122). The tissue homogenate was centrifuged for 5 min at 75 x g to remove undigested tissue. Supernatant was collected and centrifuged for 5 min at 350 x g. After centrifugation, cells were resuspended in the same medium, passed through a 40 µm cell strainer and pre-plated for 4 hours to allow for fibroblasts adherence. Non-attached cells were collected, centrifuged for 5 min at 350 x g and plated at a density of 220,000 cells/ml in IMDM with Glutamax 20% FBS, 1% Pen/Strep, 1% chicken embryo extract. Cells were seeded in µ-dish 35 mm dishes (Ibidi; 80136) pre-coated with 1% Matrigel (Corning Inc. #356231). After 3 days of proliferation, medium was replaced to induce differentiation with IMDM with Glutamax, 2% horse serum (HyClone #HYCLSH30074.02) and 1% Pen/Strep. The following day, a coating of 50% Matrigel was placed on top of the seeded cells. After upper coating, the medium was supplemented with 10 µg/ml recombinant agrin (R&D Systems #550-AG-100) and Fibroblast Growth Factor (Sigma-Aldrich #F3133).

### Drug treatments

To depolymerize microtubules, 10 µg/mL nocodazole (Sigma-Aldrich #M1404-2MG) was added to cells 5 min before live imaging acquisition.

### Plasmids and transfections

*In vitro* myofibers were transfected, 3 days after plating, with 1μg per ibidi-35mm-dish using Lipofectamine 2000 (ThermoFischer Scientific #11668-019) following manufacturer instructions. mit-dendra2 (AddGene Plasmid #55796) was used to label photoconvertible mitochondria and 4xmts-mNeonGreen (AddGene Plasmid #98876) was used to label mitochondria for immunostaining.

### Live imaging and photo-conversion

<u>Live imaging in *Drosophila*:</u> pupae were mounted for *in vivo* imaging according to the published protocol^48^ and imaged with a spinning disc confocal microscope (Olympus). Briefly, scissors were used to cut a small window in the pupal case facing one side of the thorax. Pupae were then placed in a custom-made microscope slide with a groove and slightly tilted sideways (about 20 degrees) such that the window in the pupal case and the flight muscles below face the microscope objective. Double-sided tape was applied to both sides of the groove to hold the coverslip. A drop of 50% glycerol was placed between the pupa and the coverslip to prevent the sample from drying out. During acquisition, samples were kept at 27°C in an incubation chamber at the microscope.

<u>Dendra2 photo-conversion in *Drosophila:*</u> photo-conversion was performed with a 405 nm laser for 30 sec on a square region of interest located in the middle of the flight muscle at 25 h, 27 h or 32 h APF. A dual-colour 488 nm / 561 nm acquisition was launched about 1 minute after the start of photo-conversion to track both the non-converted and the converted Dendra2 signals each 2 minutes for 30 minutes.

<u>Dendra2 photoconversion in mammalian cell culture:</u> Photoconversion experiments were performed on a Zeiss LSM 980 confocal microscope equipped with the ZEN gray edition software, using a Plan-Apochromat 63x Oil DIC objective (Numerical aperture: 1.40). Live imaging was performed using a large cage incubator to maintain cultures at 37°C and 5% CO_2_. Fluorescence photoconversion was conducted using a 405 nm wavelength laser at 100% laser power for one iteration. The photoconverted area consisted of a rectangle with 15 µm of width. Each fiber was imaged during 30 min periods at 2 min intervals, with one frame being acquired before photoconversion.

### Immunostaining of Drosophila flight muscle and image processing

Flight muscles were dissected at the indicated stage of development. Pupae were freed from their pupal case with forceps and pierced 3 times in the abdomen with a dissection needle to punch holes. Pupae were then fixed in 4% PFA in PBS + 0.3% Triton (PBST) for 30 min at room-temperature under agitation. Each fixed pupa was then dissected into two hemi-thoraxes according to a published protocol^49^. The hemi-thoraxes were incubated 2 hours in 24-well plates for actin staining with phalloidin rhodamine (1/500, Invitrogen R415) or GFP nanobody (1/2000, gift of Dirk Görlich) to visualise mitochondria and microtubules that were labelled with *Mef2*-GAL4, UAS-mit-GFP or *βTub60D*-GFP fosmid.

In pupae from the shine-GAL4 experiments the mitochondria were visualised by a two-step immunostaining. After fixation and dissection, which was done as noted above, blocking was performed for 1h RT in 5% normal goat serum (NGS) in PBS. Samples were then incubated in anti-ATP5α antibody (1/500, Abcam #ab14748) overnight, followed by secondary antibody (goat anti-mouse Alexa Fluor 488) for 2 h at room temperature. Between each step, the pupae were washed 3 times for 5 min in PBST. The hemi-thoraxes were mounted in VectaShield containing DAPI using two coverslips or sticky tape as spacers. Images from fixed flight muscles were acquired with a Zeiss LSM880 confocal with or without AiryScan super-resolution and processed with Fiji (ImageJ)^50^.

### Quantification of the photoconverted Dendra2 signal spread

A 100 µm long line was drawn from the photoconverted area, extending along the long and short axes. The plot profiles of the lines were obtained for each time point. Intensity values of the photoconverted Dendra2 signal for each position along the line were normalised to the average intensity for the corresponding time point. Normalised values greater than 1-fold were considered as true signals. The front of the signal spread was determined such that it delimited values higher and lower than 1-fold. The shift of the distance of the signal front at t=0 minutes and t=20 minutes was measured (one value per animal).

### Quantification of myofibrils width

Myofibers developing at 90h APF from *Mef2*-GAL4 or *Mef2*-Gal4, UAS-*khc-IR* flies are imaged longitudinally. Stacks are used to reconstruct a virtual cross section. Circles containing each visible cross cut myofibril were manually generated and the diameter of each circle was measured to quantify the mean myofibril diameter and its variability (standard deviation) for each animal.

### Immunostaining of in vitro cultures of myofibers

Myofibers were fixed in 4% paraformaldehyde in PBS for 10 minutes at room temperature and permeabilized with 0.5% Triton X-100 for 5 minutes. Cells were blocked in a solution containing 10% goat serum, 5% BSA and MOM block (R&D Systems). Primary and secondary antibodies were diluted unblocking solution and 0.1% saponin. Cells were incubated with primary antibodies overnight at 4°C and washed three times for 10 minutes with PBS. Cells were incubated with secondary antibodies for 1 hour at room temperature and washed three times for 10 minutes with PBS. Cells were mounted with Fluoromount-G (Invitrogen). Antibodies used for immunostaining were anti-tyrosinated α-tubulin (YL1/2) (1/25 ECACC) and anti-rat Alexa Fluor 555 secondary antibody (1/400, Life Technologies). Phalloidin conjugated with and Alexa Fluor 647 A22287(1/200, Life Technologies) was used to stain actin. DAPI was used to stain the nucleus (1/10,000; Sigma-Aldrich).

Image acquisition was performed on a Zeiss LSM 980 confocal microscope equipped with the ZEN gray edition software, using a Plan-Apochromat 63x Oil DIC objective (Numerical aperture: 1.40). All images were processed in Fiji^50^. For nocodazole treated myofibers, to account for cell drift, the Fast4Dreg Fiji plugin drift correction tool was used^51^. Graphpad Prism version 8.0.0 (San Diego, CA, USA) was used for data fitting. Data analysis was performed in Microsoft Excel and Graphpad Prism.

## Acknowledgements

We thank Manos Mavrakis for generously sharing the UAS-LAD9-GFP line before publication. We are grateful to Dirk Görlich, Yohanns Bellaïche, Thomas Rival, Ferenc Jancovics, the Vienna Drosophila Stock Centre (VDRC) and the Bloomington *Drosophila* Stock Center for fly strains and nanobodies. We are grateful to Christophe Pitaval for excellent technical support and the entire Schnorrer and Gomes labs for insightful discussions on this manuscript. We are indebted to the IBDM imaging and fly facilities for help with image acquisition, maintenance of the microscopes and production of fly food. We thank the Bioimaging and Rodent facilities of IMM for the technical support.

## Supplementary figure and video legends

**Figure S1: Individual mitochondria move at high speed.**

Stills from a high frame rate acquisition (Video S2) of a developing flight muscle fiber visualises individual mitochondria. Example mitochondria are highlighted by coloured arrows, with their initial position marked by a darker hue of the same colour. Scale bar represents 5 µm.

**Figure S2: Microtubules transiently assemble in the vicinity of myofibrils.**

**(A)** Single frames from a live acquisition (Video S6) of a developing flight muscle labelled for actin (Lifeact-Ruby, driven by *Mef2-*GAL4) and for microtubules (*Mhc*-Tau-GFP). **(B)** Flight muscles at 48 h, 72 h and 92 h APF expressing *βTub60D-*GFP and co-stained for actin (phalloidin). Note how the absence of βTub60D-GFP at 72 h and 90 h APF. Scale bars represent 10 µm in A and 5 µm in B.

**Figure S3: Microtubules are largely dispensable for myofibril maturation.**

**(A)** Scheme of temporal Spastin expression via light-activation by the shineGAL4 driver and the indicated sample collection time points. **(B, C)** Flight muscles of shine-GAL4; UAS*-spastin* pupae either raised in the dark (control, top rows) or exposed to light after 30 h APF (bottom rows) and stained for actin (phalloidin) and mitochondria (ATP5α antibody) at 48 h APF (B) or 90 h APF (C). Scale bars represent 5 µm.

**Figure S4: Kinesin dependent mitochondria localisation.**

**(A, B)** Flight muscles from *Mef2*-GAL4 control (A) and *Mef2*-GAL4, UAS-*kinesin* RNAi (*khc-IR*) (B) 90 h APF pupae were stained for actin (phalloidin) and mitochondria (mit-GFP). Note the large mitochondria aggregates outside the myofibril bundles and the thickened myofibrils in *khc-IR* pupae. **(C, D)** Quantification of sarcomere widths (C) from flight muscle of *Mef2*-GAL4 controls or *Mef2*-GAL4, UAS-*kinesin* RNAi (*khc-IR*) at 90 h APF and their standard deviations within a single animal (D). Scale bars represent 10 µm.

**Figure S5: Classification of differentiating myofibers *in vitro*.**

Classification table listing the criteria used to classify the type of mouse myofibre. Localisation and morphology of actin, mitochondria and nuclei were used to classify each myofibre into 4 distinct types, reflective of progressive stages of differentiation.

**Video S1: Mitochondria and actin morphogenesis during flight muscle development.**

Live video of developing flight muscles labelled for mitochondria (in green) and actin (in magenta) with UAS-mit-Kate2 and UAS-LAD9-GFP from 24 h APF to 31 h APF. Scale bar represents 10 µm.

**Video S2: Individual mitochondria move at high speed.**

High frame rate acquisition of a developing flight muscle fibre, in which individual mitochondria can be distinguished by expression of mit-GFP via *Mef2-*GAL4. Frames were acquired every 500 ms and the scale bar represents 5 µm.

**Video S3: Mitochondria move short-range at 25 h APF and long-range at 32 h APF.**

Live video of photo-converted mitochondria (labelled with *Mef2*-GAL4, UAS-mit-Dendra2) imaging the spread along the flight muscle long axis at 25 h APF prior to mitochondria intercalation or at 32 h APF after mitochondria intercalation. Scale bar represents 10 µm.

**Video S4: Mitochondria move long range along long axis at 32 h APF.**

Live video of photo-converted mitochondria (labelled with *Mef2*-GAL4, UAS-mit-Dendra2) imaging the spread along both, the long and the short axis of the flight muscle at 25 h and 32 h APF, prior to and after, respectively, mitochondria intercalation. Scale bar represents 10 µm.

**Video S5: Tracking single mitochondria intercalating at 27h APF.**

Live video of photo-converted mitochondria (labelled with *Mef2*-GAL4, UAS-mit-Dendra2) at 27 h APF in developing flight muscle. Note how one mitochondrion is transported away from others to rapidly intercalate into neighbouring actin bundles. Scale bar represents 5 µm.

**Video S6: Actin and microtubules during flight muscle development.**

Live video of developing flight muscle labelled for actin (in magenta) and microtubules (in green) with *Mhc*-tau-GFP and *Mef2*-GAL4, UAS-Lifeact-Ruby from 22 h APF to 29 h APF. Scale bar represents 10 µm.

## References

1. Bleck, C.K.E., Kim, Y., Willingham, T.B., and Glancy, B. (2018). Subcellular connectomic analyses of energy networks in striated muscle. Nat. Commun. 9, 5111. 10.1038/s41467-018-07676-y.

2. Katti, P., Hall, A.S., Parry, H.A., Ajayi, P.T., Kim, Y., Willingham, T.B., Bleck, C.K.E., Wen, H., and Glancy, B. (2022). Mitochondrial network configuration influences sarcomere and myosin filament structure in striated muscles. Nat. Commun. 13, 6058. 10.1038/s41467-022-33678-y.

3. Schiaffino, S. (2010). Fibre types in skeletal muscle: a personal account: Fibre types in skeletal muscle. Acta Physiol. 199, 451–463. 10.1111/j.1748-1716.2010.02130.x.

4. Spletter, M.L., and Schnorrer, F. (2014). Transcriptional regulation and alternative splicing cooperate in muscle fiber-type specification in flies and mammals. Exp. Cell Res. 321, 90–98. 10.1016/j.yexcr.2013.10.007.

5. Glancy, B., Hartnell, L.M., Malide, D., Yu, Z.-X., Combs, C.A., Connelly, P.S., Subramaniam, S., and Balaban, R.S. (2015). Mitochondrial reticulum for cellular energy distribution in muscle. Nature 523, 617–620. 10.1038/nature14614.

6. Vincent, A.E., White, K., Davey, T., Philips, J., Ogden, R.T., Lawless, C., Warren, C., Hall, M.G., Ng, Y.S., Falkous, G., et al. (2019). Quantitative 3D Mapping of the Human Skeletal Muscle Mitochondrial Network. Cell Rep. 26, 996–1009.e4. 10.1016/j.celrep.2019.01.010.

7. Avellaneda, J., Rodier, C., Daian, F., Brouilly, N., Rival, T., Luis, N.M., and Schnorrer, F. (2021). Myofibril and mitochondria morphogenesis are coordinated by a mechanical feedback mechanism in muscle. Nat. Commun. 12, 2091. 10.1038/s41467-021-22058-7.

8. Katti, P., Ajayi, P.T., Aponte, A., Bleck, C.K.E., and Glancy, B. (2022). Identification of evolutionarily conserved regulators of muscle mitochondrial network organization. Nat. Commun. 13, 6622. 10.1038/s41467-022-34445-9.

9. Ajayi, P.T., Katti, P., Zhang, Y., Willingham, T.B., Sun, Y., Bleck, C.K.E., and Glancy, B. (2022). Regulation of the evolutionarily conserved muscle myofibrillar matrix by cell type dependent and independent mechanisms. Nat. Commun. 13, 2661. 10.1038/s41467-022-30401-9.

10. Schönbauer, C., Distler, J., Jährling, N., Radolf, M., Dodt, H.-U., Frasch, M., and Schnorrer, F. (2011). Spalt mediates an evolutionarily conserved switch to fibrillar muscle fate in insects. Nature 479, 406–409. 10.1038/nature10559.

11. Gunage, R.D., Dhanyasi, N., Reichert, H., and VijayRaghavan, K. (2017). Drosophila adult muscle development and regeneration. Semin. Cell Dev. Biol. 72, 56–66. 10.1016/j.semcdb.2017.11.017.

12. Luis, N.M., and Schnorrer, F. (2021). Mechanobiology of muscle and myofibril morphogenesis. Cells Dev., 203760. 10.1016/j.cdev.2021.203760.

13. Weitkunat, M., Kaya-Çopur, A., Grill, S.W., and Schnorrer, F. (2014). Tension and Force-Resistant Attachment Are Essential for Myofibrillogenesis in Drosophila Flight Muscle. Curr. Biol. 24, 705–716. 10.1016/j.cub.2014.02.032.

14. Spletter, M.L., Barz, C., Yeroslaviz, A., Zhang, X., Lemke, S.B., Bonnard, A., Brunner, E., Cardone, G., Basler, K., Habermann, B.H., et al. (2018). A transcriptomics resource reveals a transcriptional transition during ordered sarcomere morphogenesis in flight muscle. eLife 7, e34058. 10.7554/eLife.34058.

15. Poliacikova, G., Barthez, M., Rival, T., Aouane, A., Luis, N.M., Richard, F., Daian, F., Brouilly, N., Schnorrer, F., Maurel-Zaffran, C., et al. (2023). M1BP is an essential transcriptional activator of oxidative metabolism during Drosophila development. Nat. Commun. 14, 3187. 10.1038/s41467-023-38986-5.

16. Akhmanova, A., and Kapitein, L.C. (2022). Mechanisms of microtubule organization in differentiated animal cells. Nat. Rev. Mol. Cell Biol. 23, 541–558. 10.1038/s41580-022-00473-y.

17. Denes, L.T., Kelley, C.P., and Wang, E.T. (2021). Microtubule-based transport is essential to distribute RNA and nascent protein in skeletal muscle. Nat. Commun. 12, 6079. 10.1038/s41467-021-26383-9.

18. Elhanany-Tamir, H., Yu, Y.V., Shnayder, M., Jain, A., Welte, M., and Volk, T. (2012). Organelle positioning in muscles requires cooperation between two KASH proteins and microtubules. J. Cell Biol. 198, 833–846. 10.1083/jcb.201204102.

19. Metzger, T., Gache, V., Xu, M., Cadot, B., Folker, E.S., Richardson, B.E., Gomes, E.R., and Baylies, M.K. (2012). MAP and kinesin-dependent nuclear positioning is required for skeletal muscle function. Nature 484, 120–124. 10.1038/nature10914.

20. Dhanyasi, N., VijayRaghavan, K., Shilo, B.-Z., and Schejter, E.D. (2020). Microtubules provide guidance cues for myofibril and sarcomere assembly and growth. Dev. Dyn. doi10.1002/dvdy.227.

21. Misgeld, T., and Schwarz, T.L. (2017). Mitostasis in Neurons: Maintaining Mitochondria in an Extended Cellular Architecture. Neuron 96, 651–666. 10.1016/j.neuron.2017.09.055.

22. Kaya-Çopur, A., Marchiano, F., Hein, M.Y., Alpern, D., Russeil, J., Luis, N.M., Mann, M., Deplancke, B., Habermann, B.H., and Schnorrer, F. (2021). The Hippo pathway controls myofibril assembly and muscle fiber growth by regulating sarcomeric gene expression. eLife 10, e63726. 10.7554/eLife.63726.

23. Kruppa, A.J., and Buss, F. (2021). Motor proteins at the mitochondria–cytoskeleton interface. J. Cell Sci. 134, jcs226084. 10.1242/jcs.226084.

24. Loison, O., Weitkunat, M., Kaya-Çopur, A., Nascimento Alves, C., Matzat, T., Spletter, M.L., Luschnig, S., Brasselet, S., Lenne, P.-F., and Schnorrer, F. (2018). Polarization-resolved microscopy reveals a muscle myosin motor-independent mechanism of molecular actin ordering during sarcomere maturation. PLOS Biol. 16, e2004718. 10.1371/journal.pbio.2004718.

25. Reedy, M.C., and Beall, C. (1993). Ultrastructure of Developing Flight Muscle in Drosophila. I. Assembly of Myofibrils. Dev. Biol. 160, 443–465. 10.1006/dbio.1993.1320.

26. Jankovics, F., and Brunner, D. (2006). Transiently Reorganized Microtubules Are Essential for Zippering during Dorsal Closure in Drosophila melanogaster. Dev. Cell 11, 375–385. 10.1016/j.devcel.2006.07.014.

27. Roll-Mecak, A., and Vale, R.D. (2005). The Drosophila Homologue of the Hereditary Spastic Paraplegia Protein, Spastin, Severs and Disassembles Microtubules. Curr. Biol. 15, 650–655. 10.1016/j.cub.2005.02.029.

28. Evans, K.J., Gomes, E.R., Reisenweber, S.M., Gundersen, G.G., and Lauring, B.P. (2005). Linking axonal degeneration to microtubule remodeling by Spastin-mediated microtubule severing. J. Cell Biol. 168, 599–606. 10.1083/jcb.200409058.

29. Di Pietro, F., Herszterg, S., Huang, A., Bosveld, F., Alexandre, C., Sancéré, L., Pelletier, S., Joudat, A., Kapoor, V., Vincent, J.-P., et al. (2021). Rapid and robust optogenetic control of gene expression in Drosophila. Dev. Cell 56, 3393–3404.e7. 10.1016/j.devcel.2021.11.016.

30. Schnorrer, F., Schönbauer, C., Langer, C.C.H., Dietzl, G., Novatchkova, M., Schernhuber, K., Fellner, M., Azaryan, A., Radolf, M., Stark, A., et al. (2010). Systematic genetic analysis of muscle morphogenesis and function in Drosophila. Nature 464, 287–291. 10.1038/nature08799.

31. Roman, W., Martins, J.P., Carvalho, F.A., Voituriez, R., Abella, J.V.G., Santos, N.C., Cadot, B., Way, M., and Gomes, E.R. (2017). Myofibril contraction and crosslinking drive nuclear movement to the periphery of skeletal muscle. Nat. Cell Biol. 19, 1189–1201. 10.1038/ncb3605.

32. Spletter, M.L., Barz, C., Yeroslaviz, A., Zhang, X., Lemke, S.B., Brunner, E., Cardone, G., Basler, K., Habermann, B.H., and Schnorrer, F. (2017). Systematic transcriptomics reveals a biphasic mode of sarcomere morphogenesis in flight muscles regulated by Spalt (Developmental Biology) 10.1101/229534.

33. Guo, X., Macleod, G.T., Wellington, A., Hu, F., Panchumarthi, S., Schoenfield, M., Marin, L., Charlton, M.P., Atwood, H.L., and Zinsmaier, K.E. (2005). The GTPase dMiro Is Required for Axonal Transport of Mitochondria to Drosophila Synapses. Neuron 47, 379–393. 10.1016/j.neuron.2005.06.027.

34. Stowers, R.S., Megeath, L.J., Górska-Andrzejak, J., Meinertzhagen, I.A., and Schwarz, T.L. (2002). Axonal Transport of Mitochondria to Synapses Depends on Milton, a Novel Drosophila Protein. Neuron 36, 1063–1077. 10.1016/S0896-6273(02)01094-2.

35. Zhang, X., Avellaneda, J., Spletter, M.L., Lemke, S., Mangeol, P., Habermann, B.H., and Schnorrer, F. (2023). A mechanically regulated liquid-liquid phase separation of the transcriptional regulator Tono instructs muscle development. bioRxiv, 2023.08.27.555003. 10.1101/2023.08.27.555003.

36. Gimpel, P., Lee, Y.L., Sobota, R.M., Calvi, A., Koullourou, V., Patel, R., Mamchaoui, K., Nédélec, F., Shackleton, S., Schmoranzer, J., et al. (2017). Nesprin-1α-Dependent Microtubule Nucleation from the Nuclear Envelope via Akap450 Is Necessary for Nuclear Positioning in Muscle Cells. Curr. Biol. 27, 2999–3009.e9. 10.1016/j.cub.2017.08.031.

37. Applewhite, D.A., Grode, K.D., Duncan, M.C., and Rogers, S.L. (2013). The actin-microtubule cross-linking activity of *Drosophila* Short stop is regulated by intramolecular inhibition. Mol. Biol. Cell 24, 2885–2893. 10.1091/mbc.e12-11-0798.

38. Strumpf, D., and Volk, T. (1998). Kakapo, a Novel Cytoskeletal-associated Protein Is Essential for the Restricted Localization of the Neuregulin-like Factor, Vein, at the Muscle–Tendon Junction Site. J. Cell Biol. 143, 1259–1270. 10.1083/jcb.143.5.1259.

39. Subramanian, A., Prokop, A., Yamamoto, M., Sugimura, K., Uemura, T., Betschinger, J., Knoblich, J.A., and Volk, T. (2003). Shortstop Recruits EB1/APC1 and Promotes Microtubule Assembly at the Muscle-Tendon Junction. Curr. Biol. 13, 1086–1095. 10.1016/S0960-9822(03)00416-0.

40. Elie, A., Prezel, E., Guérin, C., Denarier, E., Ramirez-Rios, S., Serre, L., Andrieux, A., Fourest-Lieuvin, A., Blanchoin, L., and Arnal, I. (2015). Tau co-organizes dynamic microtubule and actin networks. Sci. Rep. 5, 9964. 10.1038/srep09964.

41. Martins, C.S., Iv, F., Suman, S.K., Panagiotou, T.C., Sidor, C., Ruso-López, M., Plancke, C.N., Omi, S., Gomes, M., Llewellyn, A., et al. (2024). Genetically encoded reporters of actin filament organization in living cells and tissues. Preprint, 10.1101/2024.04.26.591305.

42. Chen, E.H., and Olson, E.N. (2001). Antisocial, an Intracellular Adaptor Protein, Is Required for Myoblast Fusion in Drosophila. Dev. Cell 1, 705–715. 10.1016/S1534-5807(01)00084-3.

43. Hatan, M., Shinder, V., Israeli, D., Schnorrer, F., and Volk, T. (2011). The *Drosophila* blood brain barrier is maintained by GPCR-dependent dynamic actin structures. J. Cell Biol. 192, 307–319. 10.1083/jcb.201007095.

44. El Fissi, N., Rojo, M., Aouane, A., Karatas, E., Poliacikova, G., David, C., Royet, J., and Rival, T. (2018). Mitofusin gain and loss of function drive pathogenesis in *Drosophila* models of CMT 2A neuropathy. EMBO Rep. 19, e45241. 10.15252/embr.201745241.

45. Sarov, M., Barz, C., Jambor, H., Hein, M.Y., Schmied, C., Suchold, D., Stender, B., Janosch, S., Kj, V.V., Krishnan, R., et al. (2016). A genome-wide resource for the analysis of protein localisation in Drosophila. eLife 5, e12068. 10.7554/eLife.12068.

46. Dietzl, G., Chen, D., Schnorrer, F., Su, K.-C., Barinova, Y., Fellner, M., Gasser, B., Kinsey, K., Oppel, S., Scheiblauer, S., et al. (2007). A genome-wide transgenic RNAi library for conditional gene inactivation in Drosophila. Nature 448, 151–156. 10.1038/nature05954.

47. Pimentel, M.R., Falcone, S., Cadot, B., and Gomes, E.R. (2017). In Vitro Differentiation of Mature Myofibers for Live Imaging. J. Vis. Exp., 55141. 10.3791/55141.

48. Lemke, S.B., and Schnorrer, F. (2018). In Vivo Imaging of Muscle-tendon Morphogenesis in Drosophila Pupae. J. Vis. Exp., 57312. 10.3791/57312.

49. Weitkunat, M., and Schnorrer, F. (2014). A guide to study Drosophila muscle biology. Methods 68, 2–14. 10.1016/j.ymeth.2014.02.037.

50. Schindelin, J., Arganda-Carreras, I., Frise, E., Kaynig, V., Longair, M., Pietzsch, T., Preibisch, S., Rueden, C., Saalfeld, S., Schmid, B., et al. (2012). Fiji: an open-source platform for biological-image analysis. Nat. Methods 9, 676–682. 10.1038/nmeth.2019.

51. Pylvänäinen, J.W., Laine, R.F., Saraiva, B.M.S., Ghimire, S., Follain, G., Henriques, R., and Jacquemet, G. (2023). Fast4DReg – fast registration of 4D microscopy datasets. J. Cell Sci. 136, jcs260728. 10.1242/jcs.260728.

