## Supplemental Figures for "Microtubules coordinate mitochondria transport with myofibril morphogenesis during muscle development"

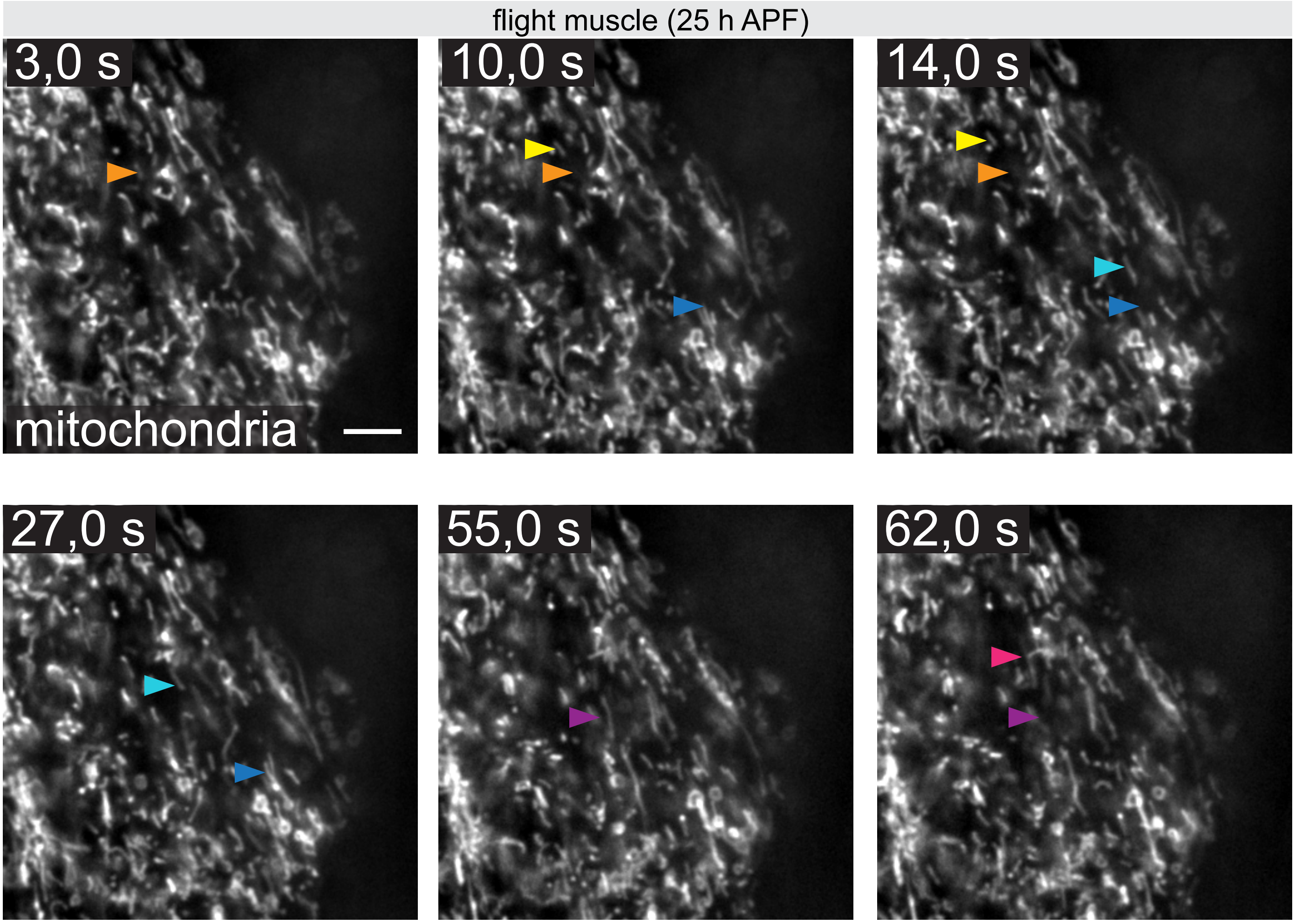


**Avellaneda et al. Figure S1**

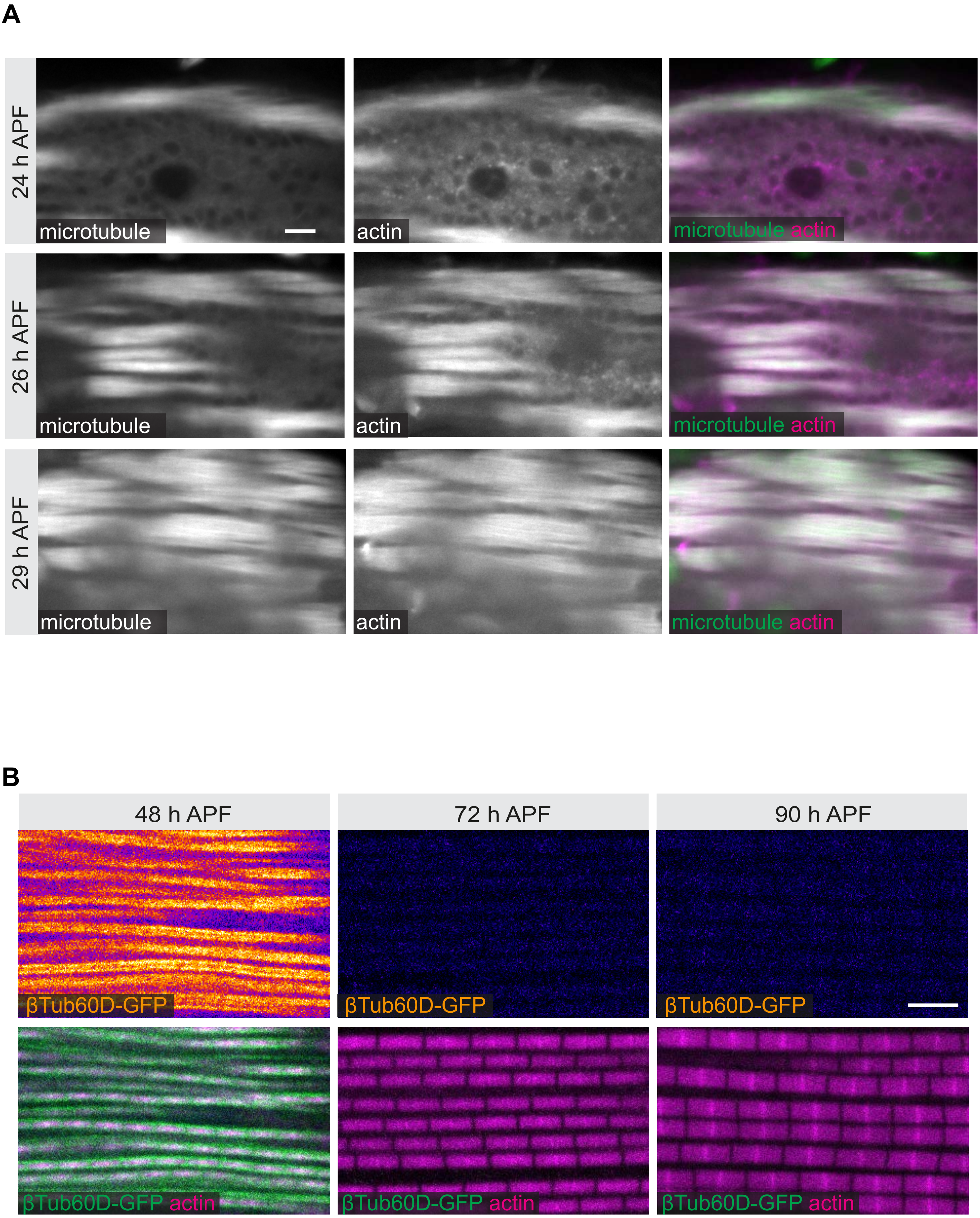


**Avellaneda et al. Figure S2**

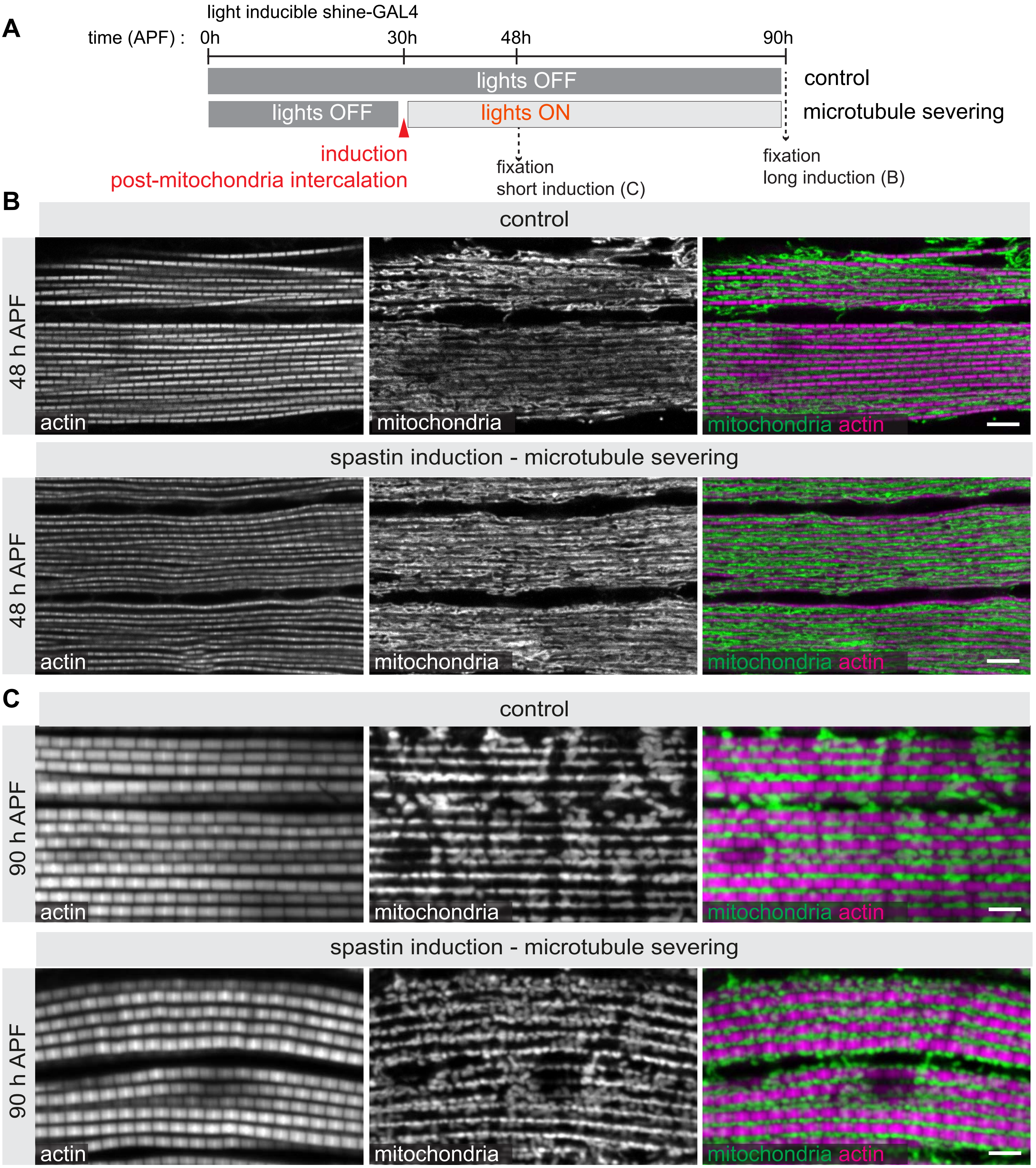


**Avellaneda et al. Figure S3**

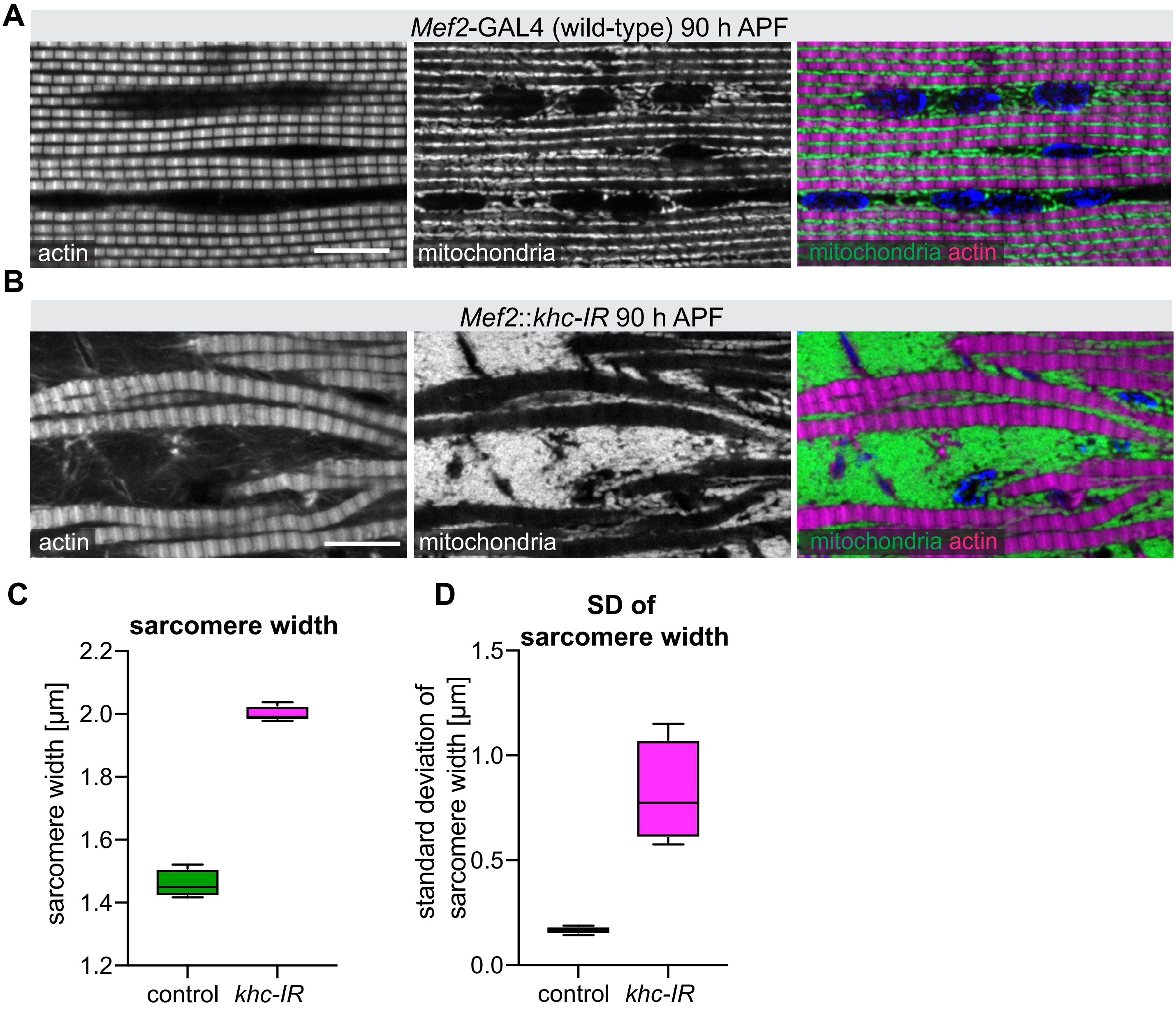


**Avellaneda et al. Figure S4**

| | Type 1 | Type 2 | Type 3 | Type 4 |
|---|---|---|---|---|
| **Nuclei** | clustered in centre of the fibre | distributed in centre of the fibre | distributed in centre of the fibre | pushed to periphery of the fibre |
| **Actin** | filaments forming a mesh at cell cortex | filaments oriented along cell axis | striations appear myofibrils visible | sarcomeres clearly defined and more mature |
| **Mitochondria** | clustered around nuclei | clustered in central parts of the fibre | in between myofibrils | squeezed between myofibrils |

**Avellaneda et al. Figure S5**
